# Drug screening in human physiologic medium identifies uric acid as an inhibitor of rigosertib efficacy

**DOI:** 10.1101/2023.07.26.550731

**Authors:** Vipin Rawat, Patrick DeLear, Prarthana Prashanth, Mete Emir Ozgurses, Anteneh Tebeje, Philippa A. Burns, Kelly O. Conger, Christopher Solís, Yasir Hasnain, Anna Novikova, Jennifer E. Endress, Paloma González-Sánchez, Wentao Dong, Greg Stephanopoulos, Gina M. DeNicola, Isaac S. Harris, David Sept, Frank M. Mason, Jonathan L. Coloff

**Author notes:** Correspondence to, 1853 W Polk St Rm 522, M/C 512, Chicago, IL 60612.

## Abstract

The non-physiological nutrient levels found in traditional culture media have been shown to affect numerous aspects of cancer cell physiology, including how cells respond to certain therapeutic agents. Here, we comprehensively evaluated how physiological nutrient levels impact therapeutic response by performing drug screening in human plasma-like medium (HPLM). We observed dramatic nutrient-dependent changes in sensitivity to a variety of FDA-approved and clinically trialed compounds, including rigosertib, an experimental cancer therapeutic that has recently failed in phase 3 clinical trials. Mechanistically, we found that the ability of rigosertib to destabilize microtubules is strongly inhibited by the purine metabolism waste product uric acid, which is uniquely abundant in humans relative to traditional *in vitro* and *in vivo* cancer models. Structural modelling studies suggest that uric acid interacts with the tubulin-rigosertib complex and may act as an uncompetitive inhibitor of rigosertib. These results offer a possible explanation for the failure of rigosertib in clinical trials and demonstrate the utility of physiological media to achieve *in vitro* results that better represent human therapeutic responses.

## INTRODUCTION

Tumor growth is influenced by both cell intrinsic and cell extrinsic factors, and nutrient availability is emerging as a critical environmental factor that can shape the metabolic fitness and proliferative capacity of tumors (1–3). Concurrent with these discoveries has been a renewed realization that standard cell culture media were not designed to mimic the nutrient environment found *in vivo*, but were rather designed to provide excess amounts of the minimal nutrients required to sustain cancer cell growth *in vitro* (4–9). As a result, the nutrients present in these traditional culture media do not accurately recapitulate the complexity or abundance of nutrients found *in vivo*. Importantly, it has recently been shown that the highly non-physiological nutrient levels found in culture media can contribute to inconsistencies between *in vitro* and *in vivo* experiments, especially for those directly related to cellular metabolism (10–13). It has also been observed that nutrient availability can affect the response to a variety of cancer therapies(14), including traditional chemotherapies (15, 16), metabolic inhibitors (17, 18), targeted therapies (19, 20), and immunomodulatory checkpoint inhibitors (21).

Because of the importance of nutrient availability in influencing tumor metabolic phenotypes and therapeutic vulnerabilities, there is considerable interest in targeting systemic metabolism either alone or in combination with existing therapies to treat cancer. This includes several dietary interventions that are being investigated as components of cancer therapies (22, 23), including amino acid starvation (24–26), ketogenic diet (27, 28), caloric restriction (29–31), and fasting-mimicking diets (32). The success of dietary intervention studies is critically dependent on mouse models and has led to tremendous interest in translating these findings to patients. And while mouse models provide an essential platform to study interactions between systemic and tumor metabolism, there are a number of metabolic differences between mice and humans that influence tumor biology (33–36), including how cancer cells respond to cancer therapeutics (15). These issues have motivated the development of novel culture media that specifically mimic the nutrient composition found in human plasma as platforms for studying therapeutic response under more physiological human nutrient conditions (12, 15).

Here, we sought to determine the extent to which nutrient availability impacts the sensitivity of cancer cells to diverse therapeutic agents by utilizing a high-throughput differential sensitivity drug-screening platform to profile therapeutic sensitivity in cancer cells growing in traditional versus physiological human plasma-like medium (HPLM). This screen revealed dramatic nutrient-dependent changes in sensitivity to a wide variety of drugs in cells cultured in HPLM. Among these differences were changes in sensitivity to the experimental therapeutic rigosertib (ON-01910), the efficacy of which was strongly antagonized by the purine degradation product uric acid. Further, our data suggests that uric acid inhibits the microtubule destabilizing function rigosertib by directly binding and weakening the rigosertib:tubulin complex. Importantly, while rigosertib has shown great promise in pre-clinical studies, it has thus far failed in phase 3 clinical trials (37, 38). Given that uric acid levels are uniquely high in humans due to the recent pseudogenization of the uricase gene (39–44), our work offers a potential explanation for the clinical failure of rigosertib. Further, our work suggests that therapies designed to lower uric acid, such as those used to treat gout and tumor lysis syndrome, may effectively augment rigosertib sensitivity in patients.

## RESULTS

### Drug screening identifies nutrient-dependent effects on therapeutic response

Commercial culture media such as DMEM and RPMI-1640 (RPMI) contain nutrients at non-physiological levels and lack many critical components present in human plasma (45). Due to these deficiencies, several labs have recently developed media that more accurately mimic physiological nutrient levels found in human circulation (10–12, 15). We have made use human plasma-like medium (HPLM) (15) to culture breast cancer cell lines, where after a two-week adaptation period we observed similar or slightly decreased growth rates (Figure S1A), and consistent remodeling of intracellular metabolite abundance (Figures S1B & S1C). Because of the recent observation of the impact of nutrient availability and cellular metabolism on the response to a variety of cancer therapies (15, 16, 20), we hypothesized that culturing in HPLM would change how cancer cells respond to therapeutic agents on a larger scale. To address this question, we utilized a high-throughput differential sensitivity drug screening platform containing a library of 626 metabolic inhibitors and anticancer compounds arrayed in 10 point dose curves (46). This platform contains compounds targeting a wide variety of cancer-relevant pathways, many of which are FDA-approved or have been evaluated in clinical trials. We screened the triple-negative breast cancer (TNBC) cell line SUM149 growing in RPMI or HPLM media, where we observed dramatic changes in sensitivity to a variety of compounds (Figure 1A and 1B, Table S1). Interestingly, while very few drugs are more effective in HPLM, a large proportion of drugs are less effective in physiological medium.

**Figure 1.**
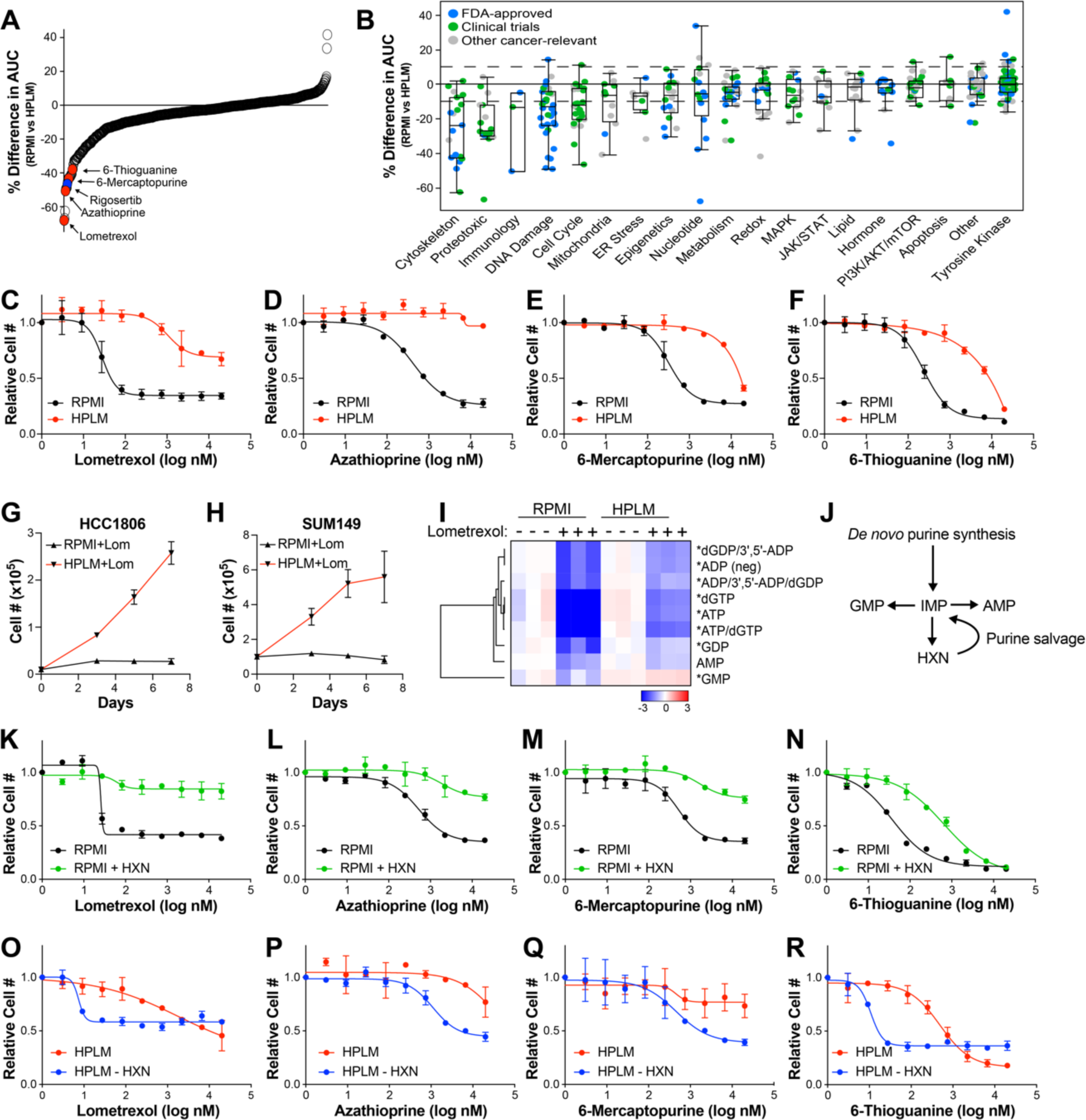
Culture in HPLM changes sensitivity to a variety of therapeutic agents. (A) Percent difference in the area under curve (% Difference in AUC) data for SUM149 cells cultured in either RPMI or HPLM after treatment with anti-cancer and metabolic inhibitor libraries. Only compounds with an Emax >50% in either medium are shown. (B) The same data as in (A) categorized based on target pathway. (C – F) Dose-response curves of the purine biosynthesis inhibitors lometrexol (C), azathioprine (D), 6-mercaptopurine (E), and 6-thioguanine (F) on SUM149 cells growing in RPMI vs HPLM. (G & H) Growth curves of HCC1806 (G) and SUM149 (H) cells treated with lometrexol in RPMI vs HPLM. (I) LC-MS analysis to quantify purine nucleotide abundance in HCC1806 cells treated with lometrexol in RPMI vs HPLM. * indicates *p* < 0.05 for HPLM + lometrexol relative to RPMI + lometrexol (unpaired two-tailed t-test). (J) Schematic representation of purine synthesis and salvage pathways. (K – N) Dose-response curves of the purine biosynthesis inhibitors lometrexol (K), azathioprine (L), 6-mercaptopurine (M), and 6-thioguanine (N) on SUM149 cells grown in RPMI with and without hypoxanthine (HXN). (O – R) Dose-response curves of the purine biosynthesis inhibitors lometrexol (O), azathioprine (P), 6-mercaptopurine (Q), and 6-thioguanine (R) on SUM149 cells grown in HPLM with and without hypoxanthine (HXN). For all panels data represents the means ± SD of triplicate samples.

Among the drugs most significantly affected by culture in HPLM are four inhibitors of the *de novo* purine biosynthesis pathway – lometrexol, azathioprine, 6-thioguanine (6-TG), and 6-mercaptopurine (6-MP) – all of which are less effective at reducing cell numbers in HPLM (Figures 1A – 1F). We found that both SUM149 cells and another TNBC cell line, HCC1806, are able to proliferate when treated with lometrexol in HPLM, but not RPMI (Figures 1G & 1H). Based on these observations, we investigated the level of purine nucleotides by LC-MS analysis in HCC1806 cells treated with and without lometrexol in both RPMI and HPLM. As expected, we found that lometrexol causes a large drop in the abundance of most purine nucleotides in RPMI; however, this drop is significantly blunted in HPLM (Figure 1I). In addition to *de novo* biosynthesis, cells can acquire purines through the purine salvage pathway (Figure 1J), and the presence of substrates for the purine salvage pathway, such as hypoxanthine, are known to reduce the efficacy of purine synthesis inhibitors (47, 48). While traditional media formulations do not contain salvage substrates, HPLM contains hypoxanthine at 10 µM as is found in human plasma. Indeed, we found that addition of hypoxanthine to RPMI is sufficient to provide resistance against these compounds (Figure 1K – 1N), and the removal of hypoxanthine from HPLM strongly increases the sensitivity of SUM149 cells to purine biosynthesis inhibitors and (Figure 1O – 1R). While the ability of hypoxanthine to rescue purine synthesis inhibitors is known, these results demonstrate the power and utility of our screening platform to identify physiological nutrients that modify cancer cell sensitivity to therapeutic agents.

### Uric acid in HPLM reduces cancer cell sensitivity to rigosertib

Another top hit from our screen was the experimental cancer therapeutic rigosertib, which is significantly less effective against cells growing in HPLM than in RPMI (Figures 1A and 2A). We validated these results by performing rigosertib dose-response analyses in HCC1806 and SUM149 cells, where we observed 2711 and 283-fold increases (respectively) in the IC50 for rigosertib in HPLM (Figures 2B and 2C). Similar results were obtained in two lung cancer cell lines, A549 and Calu6, suggesting that this effect is likely general and not restricted to breast cancer cells (Figures 2D, 2E). Rigosertib’s anti-cancer effects have been shown to be mediated by induction of both G2/M cell cycle arrest and cell death (49, 50). Accordingly, we observed that rigosertib treatment induces phosphorylation of histone H3, G2/M cell cycle arrest, and induction of cell death in RPMI but not in HPLM (Figures 2F – 2I).

**Figure 2.**
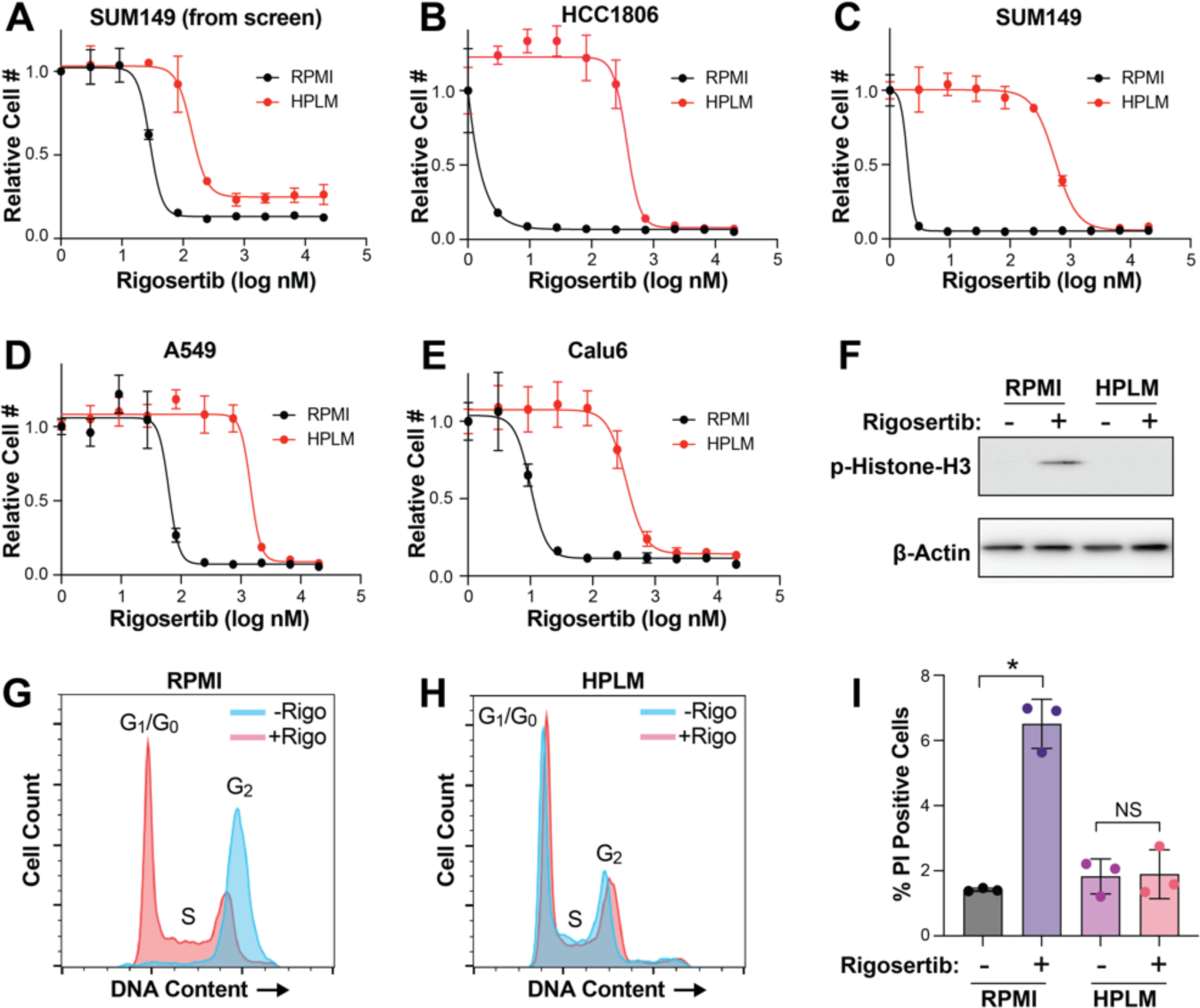
Culture in HPLM reduces sensitivity to rigosertib. (A) Dose response curve of SUM149 cells treated with rigosertib from the high-throughput screen described in Figure 1. Data are the mean ± SD of triplicate samples. (B – E) Dose response curves for rigosertib treatment of HCC1806 (B), SUM149 (C), A549 (D) and Calu6 (E) cells growing in RPMI vs HPLM. Data are the mean ± SD of triplicate samples. (F) Representative western blot of phospho-Histone H3 in HCC1806 cells treated with rigosertib in RPMI vs HPLM. (G & H) Cell cycle analysis of HCC1806 cells treated with 150 nM commercial-grade rigosertib in RPMI (G) and HPLM (H). (I) Cell death analysis of HCC1806 cells treated with 200 nM commercial-grade rigosertib in RPMI vs HPLM. Cell death and cell cycle data are the means ± SD of triplicate samples. * indicates *p* < 0.05 from unpaired two-tailed t-test. NS (not significant) indicates *p* > 0.05.

Next, we sought to determine the component(s) of HPLM that antagonizes rigosertib activity. Like RPMI and other traditional media, HPLM consists of glucose, amino acids, and salts, albeit at different concentrations (15). HPLM also contains 27 additional components that are not found in RPMI but are found in human plasma. Most of these unique ingredients are organized into 11 stock solutions numbered 8 through 18. To determine whether a unique component of HPLM is responsible for the reduced sensitivity to rigosertib, we combined HPLM stocks 8-18 and added them to RPMI and performed dose-curve analyses, where we found that stocks 8-18 were able to recapitulate the effect of HPLM on rigosertib sensitivity (Figures 3A and 3B). We then analyzed stocks 8-18 individually and found that addition of stock 18 alone was sufficient to protect against rigosertib in RPMI (Figure 3C and 3D). Importantly, stock 18 contains only one component: the purine metabolism waste product uric acid, which is present in human plasma and HPLM at 350 µM. Indeed, we found that removal of uric acid from HPLM was sufficient to dramatically increase sensitivity to rigosertib (Figures 3E). We confirmed the broad protective effects of uric acid by performing rigosertib dose curves on multiple cancer cell lines of different origin, including lung, renal and CML, where we observed the protective effects of uric acid in all cases (Figure S2). To determine whether uric acid protect cells from rigosertib in a dose-dependent manner, we performed a dose curve of uric acid in RPMI in the presence of 80 nM rigosertib. Interestingly, we found that uric acid concentrations as low as 27 µM are able to partially protect against rigosertib (Figures 3F and 3G). Similar to HPLM, uric acid alone was sufficient to block rigosertib-mediated phosphorylation of histone H3, G2/M cell cycle arrest, and cell death (Figures 3H – 3K).

**Figure 3.**
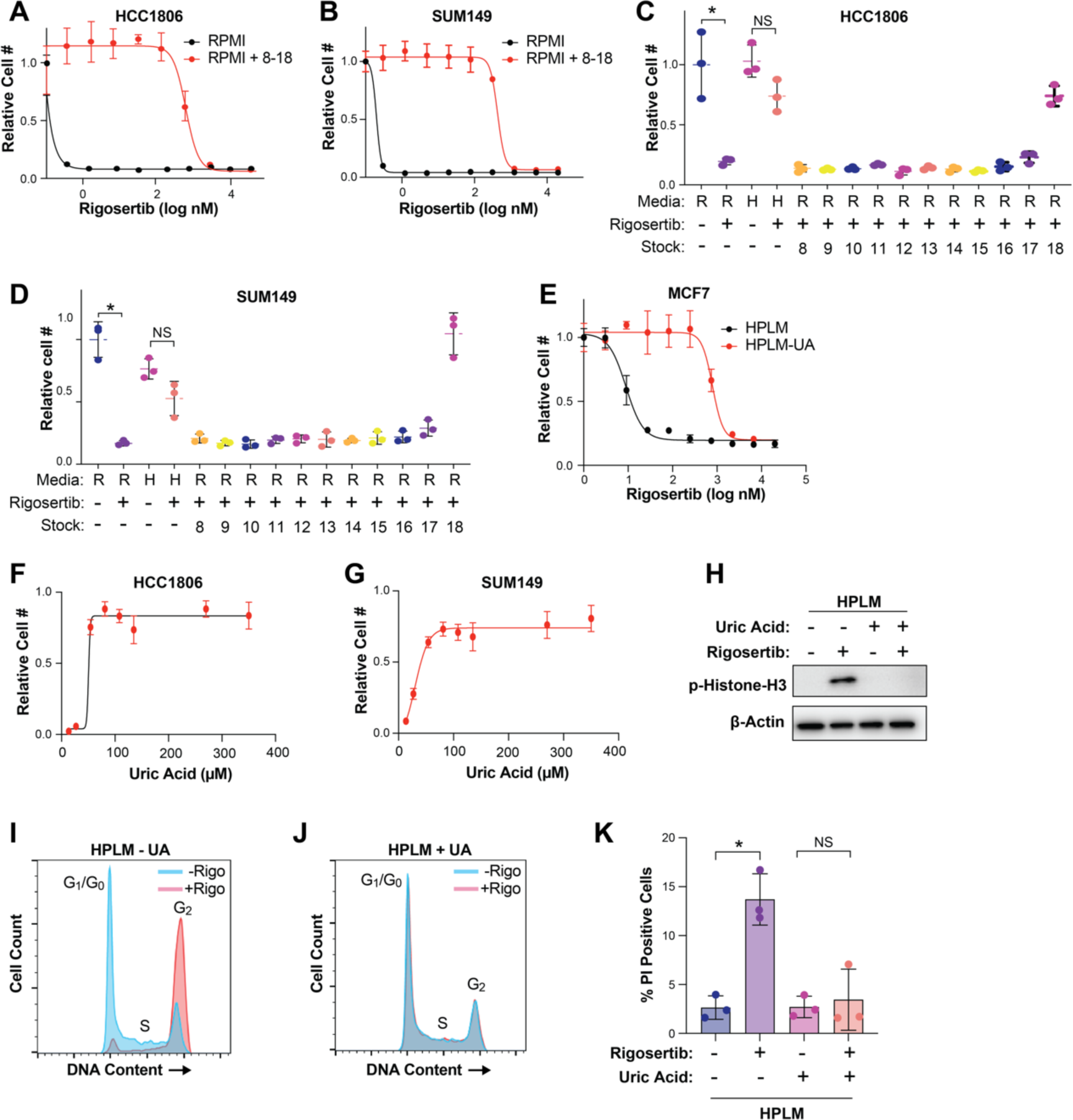
Uric acid prevents the activity of rigosertib. (A & B) Dose response curves of HCC1806 (A) and SUM149 (B) cells treated with rigosertib in RPMI vs RPMI + HPLM stocks 8-18. (C & D) Cell growth assays of HCC1806 (C) and SUM149 (D) cells treated with 80 nM rigosertib in the presence of individual HPLM stocks 8-18. R = RPMI and H = HPLM. (E) Dose response curve of MCF7 cells treated with rigosertib in HPLM vs HPLM – UA. (F & G) Dose response curves of uric acid on HCC1806 (F) and SUM149 (G) cells treated with 80 nM rigosertib. (H) Representative western blot of phospho-Histone H3 in HCC1806 cells treated with 150 nM rigosertib in HPLM vs HPLM – UA. (I & J) Cell cycle analysis of HCC1806 cells treated with 150 nM commercial-grade rigosertib in HPLM (I) and HPLM – UA (J). (K) Cell death analysis of HCC1806 cells treated with 200 nM commercial-grade rigosertib in HPLM and HPLM – UA. For all panels, data is represented as mean ± SD of triplicate samples. * indicates *p* < 0.05 from unpaired two-tailed t-test. NS (not significant) indicates *p* > 0.05.

### Uric acid inhibits the microtubule destabilizing activity of pharmaceutical grade rigosertib

While the mechanism of action of rigosertib remains controversial (51–54), several recent reports have demonstrated that rigosertib is a microtubule destabilizing agent that binds tubulin dimers at the colchicine binding site (49, 55, 56). To verify the ability of rigosertib to destabilize microtubules, we performed short-term (4 hr) treatments of HCC1806 and SUM149 cells cultured in RPMI with increasing doses of commercial-grade rigosertib, where we observed increased levels of α-tubulin in the soluble fraction of cell lysates, suggesting that rigosertib does indeed destabilize microtubules (Figures 4A, 4B, S3A, and S3B). Importantly, however, the ability of rigosertib to destabilize microtubules in cells grown in HPLM is strongly inhibited (Figures 4A, 4B, S3A, and S3B). To determine whether uric acid prevents rigosertib-mediated microtubule destabilization, we treated cells cultured in HPLM with and without 350 µM uric acid with commercial-grade rigosertib, which results in a dose-dependent increase in the level of soluble α-tubulin only in the absence of uric acid (Figures 4C, 4D, S3C and S3D).

**Figure 4.**
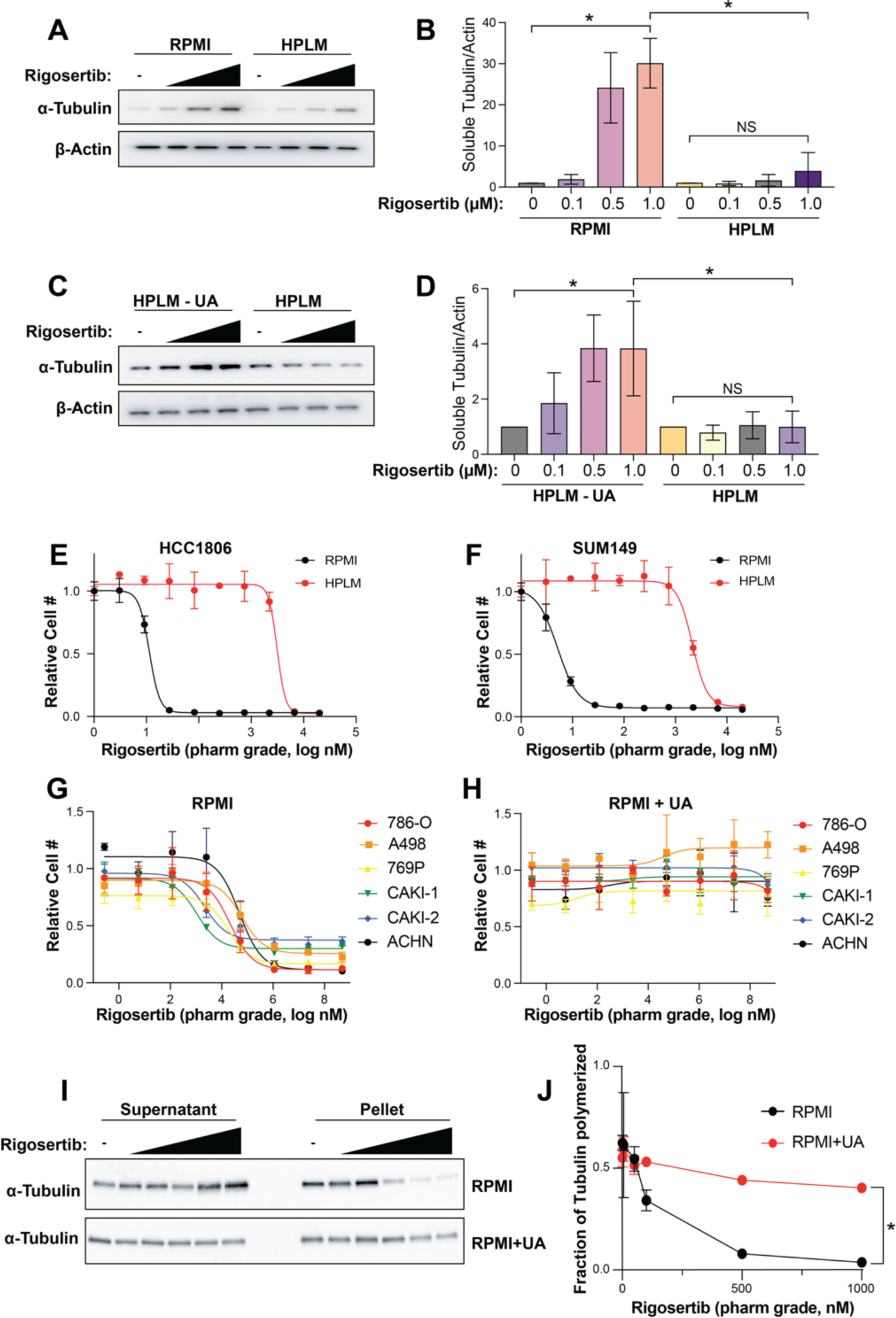
Uric acid inhibits the microtubule destabilizing activity rigosertib. (A) Western blot of soluble α-tubulin from SUM149 treated with increasing doses of rigosertib (0.1 µM, 0.5 µM and 1 µM) for 4h in RPMI and HPLM. (B) Quantification of western blots from (A). Data is represented as mean ± SD from three independent experiments. * indicates *p* < 0.05 from one way ANOVA followed by Tukey’s multiple comparison test. NS (not significant) indicates *p* > 0.05. (C) Western blot of soluble α-tubulin from SUM149 treated with increasing doses of rigosertib (0.1 µM, 0.5 µM and 1 µM) for 4h in HPLM and HPLM – UA. (D) Quantification of western blots from (C). Data is represented as mean ± SD from three independent experiments. * indicates *p* < 0.05 from one way ANOVA followed by Tukey’s multiple comparison test. NS (not significant) indicates *p* > 0.05. (E & F) Dose response curves of HCC1806 (E) and SUM149 (F) cells treated with pharmaceutical-grade rigosertib in RPMI vs HPLM. (G & H) Dose response curves of a panel of renal cancer cell lines treated with pharmaceutical-grade rigosertib in RPMI (G) vs RPMI + UA (H). (I) Western blot of soluble and pellet α-tubulin from 786-O cells treated with increasing doses (5 nM, 50 nM, 100 nM, 500 nM, 1000 nM) of pharmaceutical-grade rigosertib for 4 hr in RPMI and RPMI + UA. (J) Quantification of western blots from (I). Data is represented as means ± SD of three independent experiments. * indicates *p* < 0.05 from two-way ANOVA.

Previous work has shown that the presence of a contaminant in commercial-grade rigosertib may contribute to its anti-cancer effects (57). To determine whether HPLM blocks the effect of rigosertib or a potential contaminant, we made use of pharmaceutical-grade rigosertib that lacks the potentially active contaminant. Similar to commercial-grade rigosertib, culture of cells in HPLM strongly reduces the cellular sensitivity to pharmaceutical-grade rigosertib (Figures 4E and 4F). Importantly, addition of 350 µM uric acid to RPMI prevented sensitivity to pharmaceutical-grade rigosertib in a panel of renal cancer cell lines (Figures 4G and 4H), indicating that uric acid is protective against rigosertib and not a contaminant found in the commercial-grade compound. Similarly, treatment of 786-O cells with pharmaceutical-grade rigosertib results in a dose dependent decrease in α-tubulin found in the pellet (microtubule fraction) when compared to total tubulin in RPMI, but the addition of uric acid to RPMI prevented rigosertib-mediated destabilization of microtubules (Figures 4I and 4J).

### Uric acid binds β-tubulin and weakens the rigosertib:β-tubulin interaction

The acute ability of uric acid to prevent rigosertib-mediated destabilization of microtubules motivated us to explore the molecular effects of rigosertib and uric acid on tubulin structure. As a benchmark, we compared rigosertib to colchicine in our analyses. We started by performing molecular dynamics (MD) simulations of colchicine-bound, rigosertib-bound and apo-tubulin (non-drug bound control). After equilibrating each structure for 0.5 μs, we then performed four independent simulations of each complex resulting in more than 6 μs of total simulation time. Using principal component analysis (PCA) to evaluate the large-scale structural differences induced by colchicine and rigosertib, we found that both colchicine and rigosertib produce a similar “kink” in the dimer that likely explains their ability to prevent microtubule polymerization (Figure S4 and Supplementary Movies 1 & 2). In addition, rigosertib induces a unique conformational change that alters the relative orientation of α and β-tubulin (Figure S4, and Supplementary Movies 1 & 2). Specifically, the colchicine-bound simulation features a persistent salt bridge formed between αR221 and βE328 that is directly adjacent to the colchicine binding site (Figure 5A). Since rigosertib alters the intradimer interface and creates a greater distance between αR221 and βE328 (Figure 5B), this salt bridge cannot be formed in rigosertib-tubulin (Figure 5A). Helix H10 in β-tubulin contains E328, and loss of this salt bridge makes H10 more dynamic and creates a pocket between H10 and strand S9 (Figure 5C). Docking studies revealed that uric acid binds within this pocket via hydrogen bonding with residues in both H10 and S9 (Figure 5C). Importantly, S9 also interacts with the carboxyl group of rigosertib. Using free energy calculations we found that the net effect of uric acid binding is to weaken the binding affinity of rigosertib to β-tubulin. This data reveals a potential mechanism by which uric acid antagonizes rigosertib activity by directly interacting with the rigosertib:β-tubulin structure. Furthermore, because the uric acid binding pocket is not present in colchicine-tubulin and uric acid only binds tubulin when rigosertib is also present, our data suggests that uric acid is potentially acting as an uncompetitive inhibitor of rigosertib.

**Figure 5.**
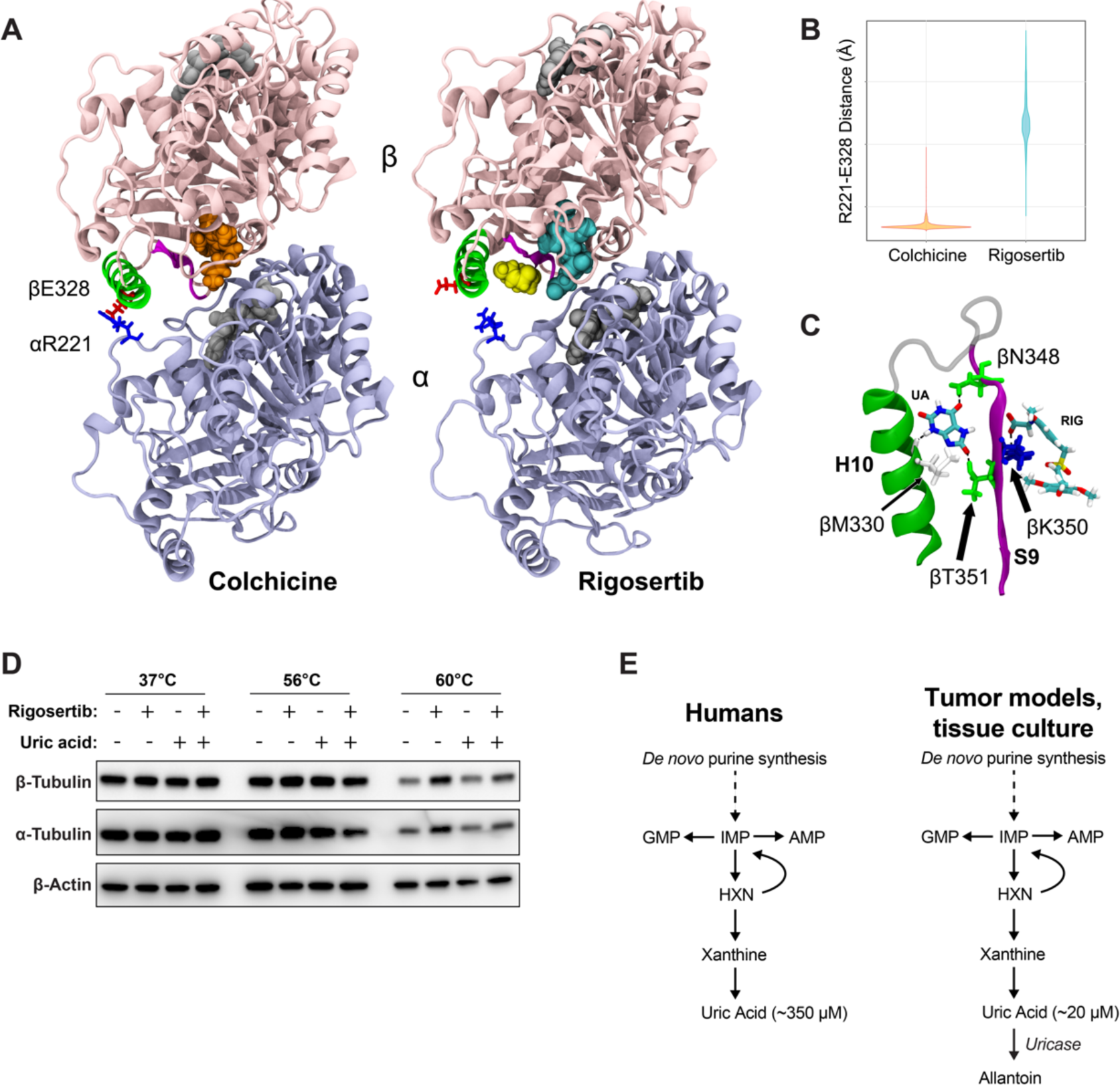
Uric acid inhibits rigosertib activity by reducing the affinity of rigosertib for β-tubulin. (A) Structural comparisons of colchicine-bound and rigosertib-bound tubulin. Colchicine and rigosertib are colored orange and cyan, respectively. The salt bridge between βE328 and αR221 found in colchicine structure is absent in the rigosertib structure, allowing H10 (green) to move away from the dimer body and create a pocket for uric acid (yellow) to bind. (B) Distance between βE328 and αR221 in the colchicine and rigosertib simulations. When this ionic bond is not formed, H10 becomes untethered which creates the binding pocket for uric acid. (C) Molecular details of uric acid binding in the pocket between H10 (green) and S9 (magenta). Residues that form hydrogen-bonds with uric acid are labeled. (D) CETSA analysis of K562 cells treated for 4 hr with 40 µM pharmaceutical-grade rigosertib in RPMI at the indicated temperature. (E) Unlike mice and other model organisms/systems, humans do not express uricase resulting in uniquely high uric acid levels.

To experimentally test whether uric acid might impair the ability of rigosertib to bind tubulin, we utilized the cellular thermal shift assay (CETSA) in cells treated with and without rigosertib and uric acid. CETSA can directly assess drug:protein interactions within cells by evaluating shifts in protein melting temperature caused by the binding of small molecules (58–60). Using this assay, we observed denaturation and precipitation of both α and β-tubulin at 60°C that was strongly reduced in the presence of rigosertib, suggesting that rigosertib is capable of binding to tubulin (Figure 5D). However, addition of uric acid to the culture media significantly reduces the stabilization of α and β-tubulin by rigosertib (Figure 5D). These results support the model suggested by our structural data that uric acid reduces the affinity of rigosertib for β-tubulin, thereby inhibiting the ability of rigosertib to destabilize microtubules.

## DISCUSSION

Despite promising pre-clinical data and extensive evaluation in early stage studies, rigosertib has thus far failed to improve outcome in the two phase 3 clinical trials in which it has been investigated (37, 38). Importantly, our discovery that uric acid strongly antagonizes the microtubule-destabilizing activity of rigosertib offers a potential explanation for the relatively poor performance of rigosertib in humans. Most species, including mice and others commonly used in cancer research (*e.g.*, bovine serum), have a functional uricase gene that converts uric acid to the more soluble allantoin, resulting in relatively low circulating uric acid concentrations (Figure 5D). However, due to the evolutionarily recent pseudogenization of the uricase gene in humans and other closely related apes, humans have circulating uric acid levels that are an order of magnitude higher than other mammals (Figure 5D) (39–44). Further, cancer patients, including those with myelodysplastic syndromes where rigosertib has been most thoroughly investigated, often present with hyperuricemia (61, 62), and therefore they may have uric acid levels that are even higher than those found in HPLM. These elevated uric acid levels could cause reduced rigosertib efficacy in humans compared to mouse models and traditional tissue culture where uric acid levels are low. Importantly, given that uric acid is the underlying cause of gout, there are numerous approved therapies to reduce uric acid levels in patients. Our work suggests that such therapies, including a low-purine diet, xanthine oxidase inhibitors (*e.g.*, allopurinol, febuxostat), and uric acid degrading enzymes (*e.g.*, rasburicase, pegloticase) could be utilized in combination with rigosertib to improve therapeutic response (62–65).

As previously mentioned, identifying the precise mechanistic target of rigosertib has been challenging. Rigosertib was initially identified as a PLK1 inhibitor (49, 66–70), and has also been proposed to act as an inhibitor of RAS (71) and PI3K (72). Use of an unbiased CRISPRi/a chemical-genetic approach combined with structural biology studies identified rigosertib as a microtubule destabilizing agent that binds to the colchicine binding site on β-tubulin (55, 56), a finding that has now been corroborated by other groups (53). Our molecular dynamic simulation studies also suggest that rigosertib binds to the colchicine binding site of β-tubulin and induces conformational changes that are similar to, but distinct from, those induced by colchicine. It is important to note, however, that the crystal structures of tubulin with colchicine or rigosertib are strongly affected by the presence of stathmin, and this limits the interpretation of these structures (55). In addition, docking results suggest that uric acid may act as an uncompetitive inhibitor of rigosertib through interaction with residues in loop S9 – a conclusion that is further confirmed by our CETSA results. Together, these results suggest that the effect of uric acid on rigosertib efficacy is mediated through microtubules and not the other proposed targets of rigosertib.

*In vitro* tissue culture models offer several advantages over *in vivo* tumor models, including the ability to perform large-scale screening studies. However, there has always been a large bottleneck of promising *in vitro* cancer findings that turn out to be irrelevant in human tumors. While there are many factors that contribute to this bottleneck, our work and that of others has shown that the non-physiological nutrient levels found in traditional culture media can contribute to some *in vitro* and *in vivo* discrepancies. Importantly, unnatural nutrient levels are not an inherent problem of tissue culture, and it is becoming clear that replacement of traditional media with more physiological media can rectify some of the problems with tissue culture systems. And while mouse models will continue to be the gold-standard of pre-clinical cancer studies, it is important to consider that there are also differences between mice and humans that can lead to discrepancies in how cancer cells respond to therapies. Our work demonstrates that use of human physiological media can lead to clinically relevant findings that otherwise might be missed using traditional tissue culture or mouse models, and we support the notion that use of physiological media is a highly valuable addition to the cancer research pipeline.

## METHODS

### Cell lines

Cell lines were acquired from the Brugge Lab at Harvard Medical School (HCC1806, SUM149), the Kim Rathmell Lab at Vanderbilt University Medical Center (A498 and Caki2), and the Jiyeon Kim (A549 and Calu6) and Vadim Gaponenko (K562) Labs at the University of Illinois at Chicago. Cell lines were tested for mycoplasma using the MycoAlert Mycoplasma Detection Kit (Lonza) and were authenticated by STR analysis. Cells were grown in human plasma-like medium according to the published formulation (15) with 5% dialyzed fetal bovine serum (FBS) (Sigma) and pen/strep (Invitrogen) at 37°C with 5% CO_2_. Media was changed at least every two days. As needed, cells were incubated in RPMI media (Thermo Fischer, 11875-093) with or without uric acid (Sigma) with 5% dialyzed FBS and pen/strep.

### Dose curve analysis

To perform dose curve analysis, 2000 cells were seeded in a 96-well plate in corresponding media. The next day cells were treated with the indicated compounds by performing nine-point serial dilutions. The media was removed after 72 hr of drug treatment and cells were fixed with 4% paraformaldehyde (Sigma F8775) for 10 min at room temperature. After fixation cells were washed with PBS and stained with 0.5 µg/ml Hoechst 33342 trihydrochloride (Thermo Fisher, H3570). Cell numbers were determined by imaging and quantifying nuclei using the Celigo imaging cytometer (Nexcelom).

### Growth curve analysis

For growth curves 10,000 cells were plated in 12-well plates and were treated with the indicated drugs the following day. Fresh media and drug were added every 2 days. After 5-7 days cells were counted using a Z1 Coulter Particle Counter (Beckman Coulter).

### Western blots

Cells were lysed in RIPA buffer (Thermo Fisher, PI89901) containing protease (Thermo Fisher, PI87786) and phosphatase inhibitor (Sigma P572, P0044) cocktails. Protein concentration was determined by BCA assay (Thermo Fisher). Quantified protein samples were separated by electrophoresis on 4–20% ready-made Tris-Glycine gels (Invitrogen) and transferred to PVDF membranes (Millipore). Membranes were blocked with 5% skim milk for 1 hr and incubated overnight with one or more primary antibodies: phospho-Histone H3 (Cell Signaling, 3377S), α-tubulin (Sigma, T9026), β-tubulin (Cell Signaling, 2146S), and β-actin (Sigma, A1978). Western blot quantification was performed using densitometry analysis on ImageJ software.

### Intracellular tubulin polymerization assay

50,000 cells per well were plated in a 12 well plate, 24 hours before treatment with increasing concentrations of rigosertib. After drug treatment, the cells were lysed in a hypotonic lysis buffer (1 mM MgCl2, 2 mM EGTA, 20 mM Tris HCl, pH 6.8, 0.15% IGEPAL, 5 µM palcitaxel) for 10 minutes at 37°C. The lysates were centrifuged at 15,000 rpm for 10 minutes at room temperature. Equal volumes of the resulting supernatants (S, containing soluble tubulin) and pellet (P, containing microtubules) for each treatment condition were subjected to SDS-PAGE followed by immunoblotting for α-tubulin.

### Cell cycle and cell death analysis

For cell cycle analysis cells were treated with 100 nM Rigosertib overnight in corresponding media. The next day cells were washed with PBS, trypsinized, quenched, and washed two times with PBS. After centrifugation, cells were fixed with ethanol at 4°C. Fixed cells were vortexed for 20 min at 4°C, washed 2 times with PBS, and stained with propidium iodide (Thermo Fisher, AAJ66584AB). Stained cells were analyzed using CytoFLEX and Gallios flow cytometers. Data was analyzed using FlowJo software. For the cell death assay cells were treated overnight with 200 nM Rigosertib in corresponding media. Trypsinized cells were suspended in 300 µl FACS buffer and stained with propidium iodide for 30 minutes. Cells were analyzed using a CytoFLEX flow cytometer and data was analyzed using FlowJo software.

### Drug screen

The MAPS platform (46) was used to test both a commercial anticancer drug library and a custom-curated metabolic inhibitor drug library. Screen was performed at the ICCB-Longwood Screening Facility (https://iccb.med.harvard.edu/small-moleculescreening). SUM149 cells were seeded at a density of 500 cells per well in a final volume of 30 µL per well of 384-well plates. After 24 hr, a Seiko Compound Transfer Robot pin transferred 100 nL of each drug library into wells with plated cells. Following pin-transfer, 20 µL of cell culture medium was added to all wells, resulting in each drug being applied at a final 10-point concentration series ranging from 20 µM to 1 nM. After 72 hr of drug treatment, the cells were washed with PBS, fixed with 4% formaldehyde, and stained with 5 mg/mL bisbenzimide. An Acumen Cellista plate cytometer was used to image plates and determine the cell numbers in individual wells. XY plots were generated comparing relative numbers of surviving RPMI and HPLM cells with concentrations of each drug tested. Area under the curve (AUC) values were calculated for each plot and drugs were ranked based on the difference between the AUCs for RPMI and HPLM cells.

### LC-MS metabolite analysis

LC-MS metabolite analysis was performed as previously described (73). Metabolites were extracted using 80% ice cold methanol. A Vanquish UPLC system was coupled to a Q Exactive HF (QE-HF) mass spectrometer equipped with HESI (Thermo Fisher). Chromatographic separation was performed with a SeQuant ZIC-pHILIC LC column, 5 μm, 150 x 4.6 mm (MilliporeSigma) with a SeQuant ZIC-pHILIC guard column, 20 x 4.6 mm (MilliporeSigma). Mobile phase A was 10 mM (NH_4_)_2_CO_3_ and 0.05% NH_4_OH in H_2_O and mobile phase B was 100% ACN. The column chamber temperature was set to 30°C. The mobile phase gradient was as follows: 0-13min: 80% to 20% of mobile phase B, 13-15min: 20% of mobile phase B. ESI ionization was performed in both positive and negative modes. The MS scan range was 60-900*m/z*. The mass resolution was 120,000 and the AGC target was 3×10^6^. The capillary voltage was 3.5 KV and the capillary temperature was 320°C. 5 μL of sample was loaded. LC-MS peaks were manually identified and integrated with EL-Maven (Elucidata) by matching with an in-house library. MetaboAnalyst was used to normalize the peak areas of target metabolites to the median fold change across all identified metabolites, calculate fold changes, and calculate p-values.

### GC-MS metabolite analysis

Polar metabolites were prepared for analysis by first drying the samples in individual microcentrifuge tubes, then adding 15 µL of methoxy amine in pyridine (MOX) (Thermo Fisher) and incubating at 40°C for 90 min. The samples were then further incubated with 20 µL of N-(tert-butyldimethylsilyl)-N-methyl-trifluoroacetamide with 1% tert-Butyldimethylchlorosilane (TBDMS) (Sigma) at 60 °C for 60 min. The resulting derivatized solution was vortexed briefly, centrifuged, and transferred into polypropylene GC/MS vials (Agilent). Subsequent metabolite abundance analysis was conducted using an Agilent 6890N GC coupled with 5975B Inert XL MS. An Agilent J&W DB-35 ms column was used. Chromatography grade helium (Airgas) was used as the carrier gas, flowing at a rate of 1 mL/min. Depending on sample abundances, either 1 or 2 µL of samples were injected using either split or splitless modes. The 6890N GC inlet temperature was set to 270 °C, and the oven temperature was initially set to 100 °C, then raised to 300 °C at a rate of 2.5 °C/min. Electron ionization mode with 70 eV was used for 5975B MS measurement. Acquisition was performed using scan mode with a detection range of 150-625 m/z. Mass isotopomer distributions (MIDs) were corrected for natural isotope abundance. Detailed methods are as published (74).

### Molecular dynamics simulations

The starting points for our simulations were the pdb structures for colchicine- and rigosertib-bound tubulin (1SA0 (75) and OV7 (55), respectively). We used a single tubulin dimer from each structure, removing the additional proteins that were added to promote crystallization. As a control, we also removed colchicine from the 1SA0 structure to create apo-tubulin to be used as a reference structure. Each complex had GTP in α-tubulin and GDP in β-tubulin, and we used CGenFF (76) to create initial force-field parameters for both colchicine and rigosertib. Each system was then solvated using TIP3 water with Na+ and Cl-added to both neutralize the system charge and set the ionic strength to 50 mM. Simulations were carried out using NAMD(77) using the CHARMM36 (78) force-field. Following heating we performed 0.5 μs equilibration simulations to remove the effects of the stathmin in the crystal structures. We then performed 4 independent 0.5 μs simulations of each structure at 300°K in an NpT ensemble with 1 atm pressure. Bonded hydrogens were fixed to allow us to use 2 fs time steps. We employed Particle Mesh Ewald (79) for long-range electrostatics and used a 10 Å cut-off and 8.5 Å switch distance for van der Waals interactions. This resulted in more than 2 μs simulation data for each of the apo, colchicine and rigosertib systems. Analysis was done using bio3D (80) and images were created using VMD (81).

### Molecular docking studies

For uric acid docking studies, we utilized Maestro (Schrödinger LLC, New York). We took a total of 40 different structures from the rigosertib and colchicine simulations (20 from each) for docking of uric acid. The following workflow was used for each structure: 1) the ligand (uric acid) and protein (tubulin-drug complexes) were prepared to be compatible with Maestro applications using LigPrep and ProteinPrep, respectively, 2) generated possible binding sites on the tubulin complex using SiteMap, 3) created receptor grids to be used for docking via Glide, and 4) docked the prepped ligand (uric acid) to the tubulin complex using ligand docking by Glide. To evaluate the binding free energy of rigosertib with and without uric acid, we utilized the Molecular Mechanics Generalized Born Surface Area methods (MMGBSA) in Prime.

### Cellular thermal shift assay (CETSA)

The K562 cells were treated with 40 µM Rigosertib for 4 hr in the corresponding media. After 4h, cells were washed with 1X PBS. Next, cells were resuspended in PBS containing 1X Halt protease inhibitor cocktail (Thermo Fisher, PI87786) and counted using a Z1 Coulter Particle Counter (Beckman Coulter). Following cell count, 700,000 cells were dispensed in PCR tube and heated at the indicated temperatures for 3 minutes in a thermocycler. After heating, cells were cooled to 20°C and lysed by thee cycles of freeze/thawi in liquid nitrogen. Following lysis, denatured proteins were separated by centrifugation at 15,000 rpm for 10 min at 4°C. The lysate was dissolved in 6X loading buffer and run on SDS-PAGE as described in the western blot section.

### Statistics

Statistical analyses were performed using GraphPad Prism9 and Microsoft Excel. Unpaired two-tailed t-test was used in most of the experiments. Where applicable one-way ANOVA followed by Tukey’s multiple comparisons test and two-way ANOVA was used.

## ACKNOWLEDGEMENTS

We would like to thank Huiping Zhao, Dan Sackett, Jiyeon Kim, and Dan Shaye for reagents, technical assistance, and/or helpful conversations. We also thanks Stephen Cosenza from Onconova Therapeutics for providing pharmaceutical-grade rigosertib. We also thank Joan Brugge and the Ludwig Center and Harvard for their support. This work made use of facilities in the Flow Cytometry Core lab (Research Resources Center, UIC) and Proteomics/Metabolomics Core facility (Moffitt Cancer Center). We would also like to thank ICCB-Longwood Screening Facility. This research was supported by the National Cancer Institute (K22 CA215828 to J.L.C. & R37 CA230042 to G.M.D.) and the Department of Defense (W81XWH2110786 to F.M.M.).

## AUTHOR CONTRIBUTIONS

Conceptualization, J.L.C. and V.R.; methodology, J.L.C., V.R., P.P., M.E.O., P.D., P.G.S, I.H., D.S.; validation, J.L.C., V.R., P.P., A.T, P.D, M.E.O.; formal analysis, J.L.C., V.R., P.P., P.A.B., K.O.C., M.E.O.; investigation, J.L.C., V.R., P.P, M.E.O, A.T., J.E.R., P.D. P.A.B., K.O.C., C.S.,Y.H., A.N., P.G.S., G.M.D., W.K.R., I.H., D.S., F.M.M, G.S., W.D.; resources, J.L.C., V.R, I.H. D.S.; data curation, J.L.C. and V.R.; writing – original draft, J.L.C., V.R., M.E.O., P.P., P.D.; writing, review & editing, J.L.C., V.R., P.P., M.E.O and P.D.; visualization, J.L.C., V.R., and.; supervision, J.L.C.; project administration, J.L.C.; funding acquisition, J.L.C.

---

**Supplementary Figure 1.**
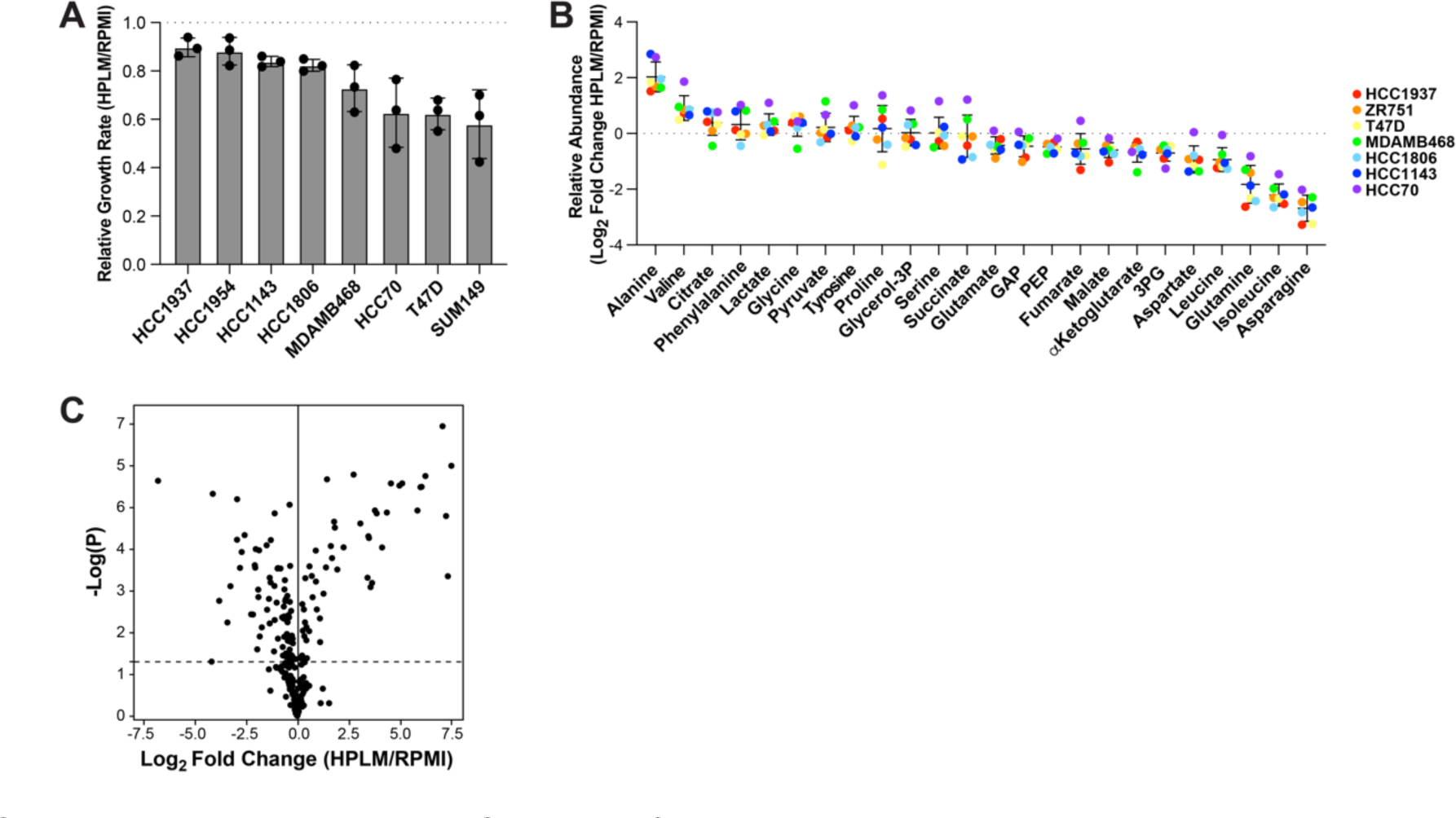
Growth of breast cancer cells in HPLM causes remodeling of intracellular metabolite abundance. Related to Figure 1. (A) Relative growth rate of breast cancer cell lines cultured in HPLM vs RPMI. Data is represented as mean ± SD from triplicate samples. (B) GC-MS analysis of relative metabolite abundance in breast cancer cells growing in HPLM vs RPMI. Data is represented as the mean of triplicate samples for each cell line. (C) LC-MS analysis of relative metabolite abundance in HCC1806 cells growing in HPLM vs RPMI. Data is represented as the mean of triplicate samples.

**Supplementary Figure 2.**
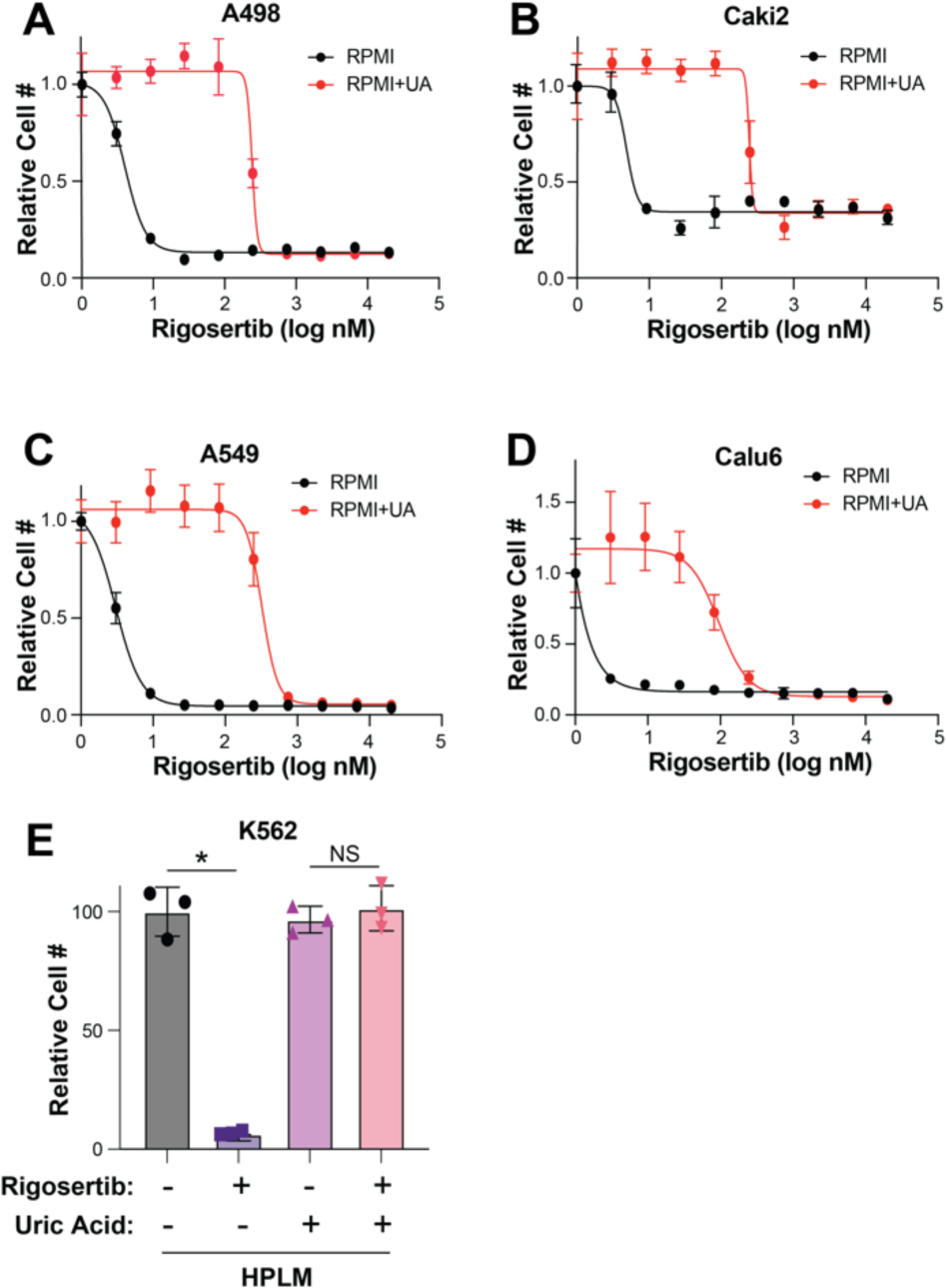
Uric acid reduces sensitivity to rigosertib in diverse cell lines. Related to Figure 3. (A – D) Dose response curves of A498 (A), Caki2 (B), A549 (C), and Calu6 (D) cells treated with rigosertib in RPMI vs RPMI + uric acid. (E) Growth of K562 cells treated with 100 nM rigosertib in HPLM – UA vs HPLM. For all panels, data is represented as mean ± SD of triplicate samples. * indicates *p* < 0.05 from unpaired two-tailed t-test. NS (not significant) indicates *p* > 0.05.

**Supplementary Figure 3.**
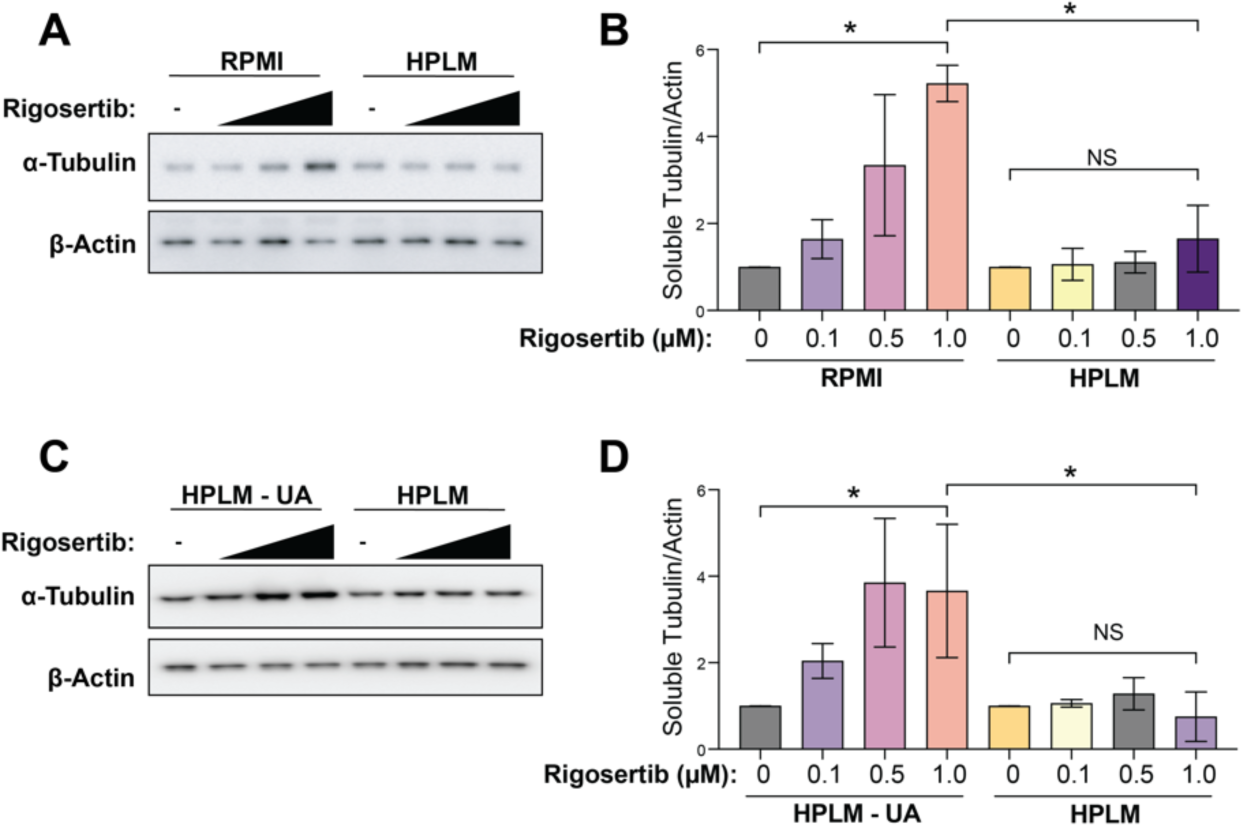
Uric acid inhibits the microtubule destabilizing activity rigosertib. Related to figure 4. (A) Western blot of soluble α-tubulin from HCC1806 treated with increasing doses of rigosertib (0.1 µM, 0.5 µM and 1 µM) for 4 hr in RPMI and HPLM. (B) Quantification of western blots from (A). Data is represented as mean ± SD from three independent experiments. * indicates *p* < 0.05 from one way ANOVA followed by Tukey’s multiple comparison test. NS (not significant) indicates *p* > 0.05. (C) Western blot of soluble α-tubulin from HCC1806 treated with increasing doses of rigosertib (0.1 µM, 0.5 µM and 1 µM) for 4 hr in HPLM and HPLM – UA. (D) Quantification of western blots from (C). Data is represented as mean ± SD from three independent experiments. * indicates *p* < 0.05 from one way ANOVA followed by Tukey’s multiple comparison test. NS (not significant) indicates *p* > 0.05.

**Supplementary Figure 4.**
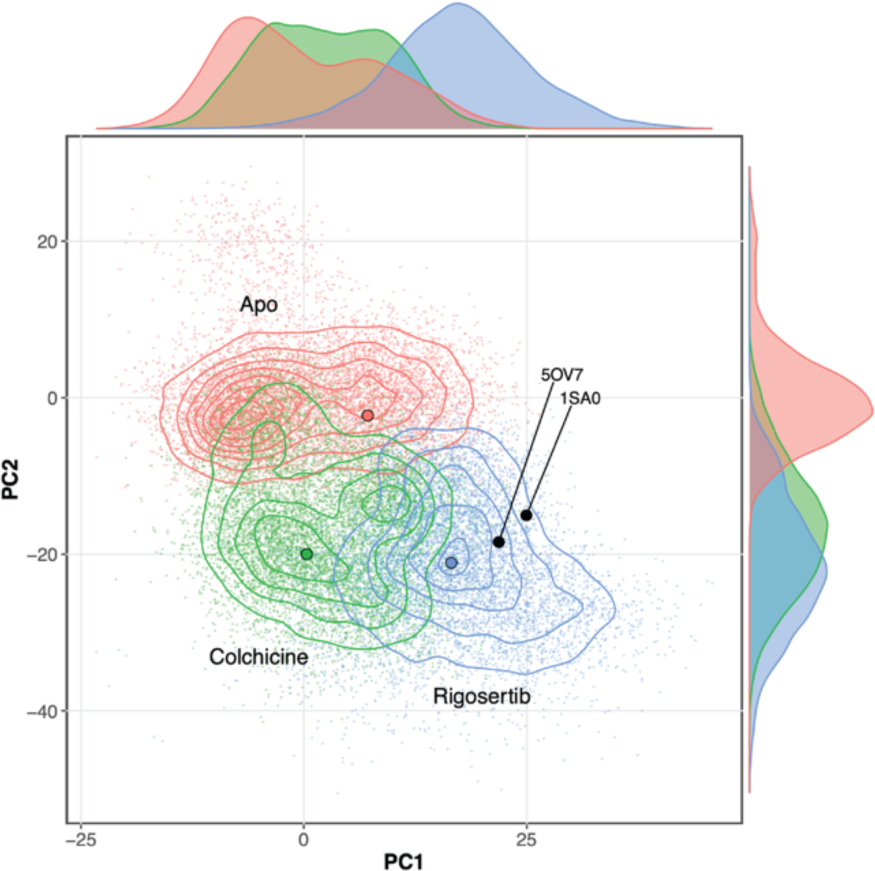
Rigosertib and colchicine have overlapping effects on tubulin structure. Comparison of apo (red), colchicine-bound (green), and rigosertib-bound tubulin (blue) using PCA analysis. Crystal structures of rigosertib (5OV7) and colchicine (1SA0) are superimposed on the PCA space, highlighting some of the limited information in the crystal structures. Both colchicine and rigosertib cause a conformational change that makes the dimer non-polymerizable (PC2), but rigosertib induces an additional conformational change that is distinct from colchicine (PC1).

## REFERENCES

1. Altea-Manzano P, et al. Nutrient metabolism and cancer in the in vivo context: a metabolic game of give and take. EMBO Rep 2020;21(10):e50635.

2. Sullivan MR, et al. Quantification of microenvironmental metabolites in murine cancers reveals determinants of tumor nutrient availability. Elife 2019;8. doi:10.7554/eLife.44235

3. Rinaldi G, Rossi M, Fendt S-M. Metabolic interactions in cancer: cellular metabolism at the interface between the microenvironment, the cancer cell phenotype and the epigenetic landscape. Wiley Interdiscip Rev Syst Biol Med 2018;10(1). doi:10.1002/wsbm.1397

4. Dulbecco R, Freeman G. Plaque production by the polyoma virus. Virology 1959;8(3):396–397.

5. Eagle H. The specific amino acid requirements of a human carcinoma cell (Stain HeLa) in tissue culture. J Exp Med 1955;102(1):37–48.

6. Eagle H. The specific amino acid requirements of a mammalian cell (strain L) in tissue culture. J Biol Chem 1955;214(2):839–852.

7. Eagle H. Nutrition needs of mammalian cells in tissue culture. Science 1955;122(3168):501–514.

8. Eagle H. Amino acid metabolism in mammalian cell cultures. Science 1959;130(3373):432–437.

9. Moore GE, Gerner RE, Franklin HA. Culture of normal human leukocytes. JAMA 1967;199(8):519–524.

10. Biancur DE, et al. Compensatory metabolic networks in pancreatic cancers upon perturbation of glutamine metabolism. Nat Commun 2017;8:15965.

11. Davidson SM, et al. Environment Impacts the Metabolic Dependencies of Ras-Driven Non-Small Cell Lung Cancer. Cell Metab 2016;23(3):517–528.

12. Vande Voorde J, et al. Improving the metabolic fidelity of cancer models with a physiological cell culture medium. Sci Adv 2019;5(1):eaau7314.

13. Barekatain Y, et al. Homozygous MTAP deletion in primary human glioblastoma is not associated with elevation of methylthioadenosine. Nat Commun 2021;12(1):4228.

14. Muir A, Vander Heiden MG. The nutrient environment affects therapy. Science 2018;360(6392):962–963.

15. Cantor JR, et al. Physiologic Medium Rewires Cellular Metabolism and Reveals Uric Acid as an Endogenous Inhibitor of UMP Synthase. Cell 2017;169(2):258–272.e17.

16. Kanarek N, et al. Histidine catabolism is a major determinant of methotrexate sensitivity. Nature 2018;559(7715):632–636.

17. Muir A, et al. Environmental cystine drives glutamine anaplerosis and sensitizes cancer cells to glutaminase inhibition. Elife 2017;6. doi:10.7554/eLife.27713

18. Gui DY, et al. Environment Dictates Dependence on Mitochondrial Complex I for NAD+ and Aspartate Production and Determines Cancer Cell Sensitivity to Metformin. Cell Metab 2016;24(5):716–727.

19. Torres-Quesada O, et al. Physiological Cell Culture Media Tune Mitochondrial Bioenergetics and Drug Sensitivity in Cancer Cell Models. Cancers (Basel) 2022;14(16). doi:10.3390/cancers14163917

20. Palm W, et al. The Utilization of Extracellular Proteins as Nutrients Is Suppressed by mTORC1. Cell 2015;162(2):259–270.

21. Kouidhi S, Ben Ayed F, Benammar Elgaaied A. Targeting Tumor Metabolism: A New Challenge to Improve Immunotherapy. Front Immunol 2018;9:353.

22. Kanarek N, Petrova B, Sabatini DM. Dietary modifications for enhanced cancer therapy. Nature 2020;579(7800):507–517.

23. Tajan M, Vousden KH. Dietary Approaches to Cancer Therapy. Cancer Cell 2020;37(6):767–785.

24. Maddocks ODK, et al. Modulating the therapeutic response of tumours to dietary serine and glycine starvation. Nature 2017;544(7650):372–376.

25. Maddocks ODK, et al. Serine starvation induces stress and p53-dependent metabolic remodelling in cancer cells. Nature 2013;493(7433):542–546.

26. Gao X, et al. Dietary methionine influences therapy in mouse cancer models and alters human metabolism. Nature 2019;572(7769):397–401.

27. Ferrere G, et al. Ketogenic diet and ketone bodies enhance the anticancer effects of PD-1 blockade. JCI Insight 2021;6(2):145207.

28. Hopkins BD, et al. Suppression of insulin feedback enhances the efficacy of PI3K inhibitors. Nature 2018;560(7719):499–503.

29. Kalaany NY, Sabatini DM. Tumours with PI3K activation are resistant to dietary restriction. Nature 2009;458(7239):725–731.

30. Mao Y-Q, et al. The antitumour effects of caloric restriction are mediated by the gut microbiome. Nat Metab 2023;5(1):96–110.

31. Lien EC, et al. Low glycaemic diets alter lipid metabolism to influence tumour growth. Nature 2021;599(7884):302–307.

32. Caffa I, et al. Fasting-mimicking diet and hormone therapy induce breast cancer regression. Nature 2020;583(7817):620–624.

33. Adelman R, Saul RL, Ames BN. Oxidative damage to DNA: relation to species metabolic rate and life span. Proc Natl Acad Sci U S A 1988;85(8):2706–2708.

34. Demetrius L. Of mice and men. When it comes to studying ageing and the means to slow it down, mice are not just small humans. EMBO Rep 2005;6 Spec No(Suppl 1):S39–44.

35. Terpstra AH. Differences between humans and mice in efficacy of the body fat lowering effect of conjugated linoleic acid: role of metabolic rate. J Nutr 2001;131(7):2067–2068.

36. Meyer MJ, et al. Differences in Metformin and Thiamine Uptake between Human and Mouse Organic Cation Transporter 1: Structural Determinants and Potential Consequences for Intrahepatic Concentrations. Drug Metab Dispos 2020;48(12):1380–1392.

37. Garcia-Manero G, et al. Rigosertib versus best supportive care for patients with high-risk myelodysplastic syndromes after failure of hypomethylating drugs (ONTIME): a randomised, controlled, phase 3 trial. Lancet Oncol 2016;17(4):496–508.

38. O’Neil BH, et al. A phase II/III randomized study to compare the efficacy and safety of rigosertib plus gemcitabine versus gemcitabine alone in patients with previously untreated metastatic pancreatic cancer. Ann Oncol 2015;26(9):1923–1929.

39. Yeldandi AV, et al. Molecular evolution of the urate oxidase-encoding gene in hominoid primates: nonsense mutations. Gene 1991;109(2):281–284.

40. Oda M, et al. Loss of urate oxidase activity in hominoids and its evolutionary implications.Mol Biol Evol 2002;19(5):640–653.

41. Wu XW, et al. Urate oxidase: primary structure and evolutionary implications. Proc Natl Acad Sci U S A 1989;86(23):9412–9416.

42. Wu XW, et al. Two independent mutational events in the loss of urate oxidase during hominoid evolution. J Mol Evol 1992;34(1):78–84.

43. Kratzer JT, et al. Evolutionary history and metabolic insights of ancient mammalian uricases. Proc Natl Acad Sci U S A 2014;111(10):3763–3768.

44. Varela-Echavarría A, Montes de Oca-Luna R, Barrera-Saldaña HA. Uricase protein sequences: conserved during vertebrate evolution but absent in humans. FASEB J 1988;2(15):3092–3096.

45. Psychogios N, et al. The human serum metabolome. PLoS One 2011;6(2):e16957.

46. Harris IS, et al. Deubiquitinases Maintain Protein Homeostasis and Survival of Cancer Cells upon Glutathione Depletion. Cell Metab 2019;29(5):1166–1181.e6.

47. Goday A, et al. Purine salvage pathway in leukemic cells. Adv Exp Med Biol 1979;122B:357–363.

48. Natsumeda Y, et al. Significance of purine salvage in circumventing the action of antimetabolites in rat hepatoma cells. Cancer Res 1989;49(1):88–92.

49. Kowalczyk JT, et al. Rigosertib Induces Mitotic Arrest and Apoptosis in RAS-Mutated Rhabdomyosarcoma and Neuroblastoma. Mol Cancer Ther 2021;20(2):307–319.

50. Hyoda T, et al. Rigosertib induces cell death of a myelodysplastic syndrome-derived cell line by DNA damage-induced G2/M arrest. Cancer Sci 2015;106(3):287–293.

51. Ritt DA, et al. Inhibition of Ras/Raf/MEK/ERK Pathway Signaling by a Stress-Induced Phospho-Regulatory Circuit. Mol Cell 2016;64(5):875–887.

52. Steegmaier M, et al. BI 2536, a potent and selective inhibitor of polo-like kinase 1, inhibits tumor growth in vivo. Curr Biol 2007;17(4):316–322.

53. Twarog NR, et al. Robust Classification of Small-Molecule Mechanism of Action Using a Minimalist High-Content Microscopy Screen and Multidimensional Phenotypic Trajectory Analysis. PLoS One 2016;11(2):e0149439.

54. Mäki-Jouppila JHE, et al. Centmitor-1, a novel acridinyl-acetohydrazide, possesses similar molecular interaction field and antimitotic cellular phenotype as rigosertib, on 01910.Na. Mol Cancer Ther 2014;13(5):1054–1066.

55. Jost M, et al. Combined CRISPRi/a-Based Chemical Genetic Screens Reveal that Rigosertib Is a Microtubule-Destabilizing Agent. Mol Cell 2017;68(1):210–223.e6.

56. Jost M, et al. Pharmaceutical-Grade Rigosertib Is a Microtubule-Destabilizing Agent. Mol Cell 2020;79(1):191–198.e3.

57. Baker SJ, et al. A Contaminant Impurity, Not Rigosertib, Is a Tubulin Binding Agent. Mol Cell 2020;79(1):180–190.e4.

58. Bisserier M, Wajapeyee N. Mechanisms of resistance to EZH2 inhibitors in diffuse large B-cell lymphomas. Blood 2018;131(19):2125–2137.

59. Langebäck A, et al. CETSA-based target engagement of taxanes as biomarkers for efficacy and resistance. Sci Rep 2019;9(1):19384.

60. Martinez Molina D, et al. Monitoring drug target engagement in cells and tissues using the cellular thermal shift assay. Science 2013;341(6141):84–87.

61. Feng Y, Jiang T, Wang L. Hyperuricemia and acute kidney injury secondary to spontaneous tumor lysis syndrome in low risk myelodysplastic syndrome. BMC Nephrol 2014;15:164.

62. Hummel M, et al. Effective treatment and prophylaxis of hyperuricemia and impaired renal function in tumor lysis syndrome with low doses of rasburicase. Eur J Haematol 2008;80(4):331–336.

63. Suzuki S, et al. Comparison between febuxostat and allopurinol uric acid-lowering therapy in patients with chronic heart failure and hyperuricemia: a multicenter randomized controlled trial. J Int Med Res 2021;49(12):3000605211062770.

64. Coiffier B, et al. Guidelines for the management of pediatric and adult tumor lysis syndrome: an evidence-based review. J Clin Oncol 2008;26(16):2767–2778.

65. Kennedy LD, Ajiboye VO. Rasburicase for the prevention and treatment of hyperuricemia in tumor lysis syndrome. J Oncol Pharm Pract 2010;16(3):205–213.

66. Gumireddy K, et al. ON01910, a non-ATP-competitive small molecule inhibitor of Plk1, is a potent anticancer agent. Cancer Cell 2005;7(3):275–286.

67. Radke K, et al. Anti-tumor effects of rigosertib in high-risk neuroblastoma. Transl Oncol 2021;14(8):101149.

68. Günther JK, et al. Rigosertib-Activated JNK1/2 Eliminate Tumor Cells through p66Shc Activation. Biology (Basel) 2020;9(5). doi:10.3390/biology9050099

69. Xu F, et al. Rigosertib as a selective anti-tumor agent can ameliorate multiple dysregulated signaling transduction pathways in high-grade myelodysplastic syndrome. Sci Rep 2014;4:7310.

70. Zhou X, et al. Efficacy of rigosertib, a small molecular RAS signaling disrupter for the treatment of KRAS-mutant colorectal cancer. Cancer Biol Med 2021;19(2):213–228.

71. Athuluri-Divakar SK, et al. A Small Molecule RAS-Mimetic Disrupts RAS Association with Effector Proteins to Block Signaling. Cell 2016;165(3):643–655.

72. Prasad A, et al. Styryl sulfonyl compounds inhibit translation of cyclin D1 in mantle cell lymphoma cells. Oncogene 2009;28(12):1518–1528.

73. Kang YP, et al. Non-canonical Glutamate-Cysteine Ligase Activity Protects against Ferroptosis. Cell Metab 2021;33(1):174–189.e7.

74. Wentao Dong, et al. Oncogenic metabolic rewiring independent of proliferative control in human mammary epithelial cells. bioRxiv 2022;2022.04.08.486845.

75. Ravelli RBG, et al. Insight into tubulin regulation from a complex with colchicine and a stathmin-like domain. Nature 2004;428(6979):198–202.

76. Vanommeslaeghe K, et al. CHARMM general force field: A force field for drug-like molecules compatible with the CHARMM all-atom additive biological force fields. J Comput Chem 2010;31(4):671–690.

77. Phillips JC, et al. Scalable molecular dynamics on CPU and GPU architectures with NAMD. J Chem Phys 2020;153(4):044130.

78. Best RB, et al. Optimization of the Additive CHARMM All-Atom Protein Force Field Targeting Improved Sampling of the Backbone ϕ, ψ and Side-Chain χ1 and χ2 Dihedral Angles. J. Chem. Theory Comput. 2012;8(9):3257–3273.

79. Essmann U, et al. A smooth particle mesh Ewald method. The Journal of Chemical Physics 1995;103(19):8577–8593.

80. Grant BJ, et al. Bio3d: an R package for the comparative analysis of protein structures. Bioinformatics 2006;22(21):2695–2696.

81. Humphrey W, Dalke A, Schulten K. VMD: visual molecular dynamics. J Mol Graph 1996;14(1):33–8, 27–28.

